# Extensive co-regulation of neighbouring genes complicates the use of eQTLs in target gene prioritisation

**DOI:** 10.1101/2023.09.29.560109

**Authors:** Ralf Tambets, Anastassia Kolde, Peep Kolberg, Michael I Love, Kaur Alasoo

## Abstract

Identifying causal genes underlying genome-wide association studies (GWAS) is a fundamental problem in human genetics. Although colocalisation with gene expression quantitative trait loci (eQTLs) is often used to prioritise GWAS target genes, systematic benchmarking has been limited due to unavailability of large ground truth datasets. Here, we re-analysed plasma protein QTL data from 3,301 individuals of the INTERVAL cohort together with 131 eQTL Catalogue datasets. Focusing on variants located within or close to the affected protein identified 793 proteins with at least one *cis*-pQTL where we could assume that the most likely causal gene was the gene coding for the protein. We then benchmarked the ability of *cis*-eQTLs to recover these causal genes by comparing three Bayesian colocalisation methods (coloc.susie, coloc.abf and CLPP) and five Mendelian randomisation (MR) approaches (three varieties of inverse-variance weighted MR, MR-RAPS, and MRLocus). We found that assigning fine-mapped pQTLs to their closest protein coding genes outperformed all colocalisation methods regarding both precision (71.9%) and recall (76.9%). Furthermore, the colocalisation method with the highest recall (coloc.susie - 46.3%) also had the lowest precision (45.1%). Combining evidence from multiple conditionally distinct colocalising QTLs with MR increased precision to 81%, but this was accompanied by a large reduction in recall to 7.1%. Furthermore, the choice of the MR method greatly affected performance, with the standard inverse-variance weighted MR often producing many false positives. Our results highlight that linking GWAS variants to target genes remains challenging with eQTL evidence alone, and prioritising novel targets requires triangulation of evidence from multiple sources.

## Introduction

Linking non-coding regulatory variants from genome-wide association studies (GWAS) to their causal target genes is a fundamental problem in human genetics. Although colocalisation with gene expression quantitative trait loci (eQTLs) is often used to assign GWAS hits to target genes, methodological differences between studies and lack of ground truth have made it challenging to systematically evaluate the performance of this approach.

Even for the simple strategy of assigning each GWAS variant to the closest gene, different studies have yielded varying estimates of precision and recall depending on which gene-variant pairs are being used as the truth set [1–4]. Among others, these studies include the locus-2-gene (L2G) model [1], the activity-by-contact (ABC) model [2], ProGeM [3], and polygenic priority score (PoPS) [4]. The L2G model used 445 “gold-standard-positive” genes selected manually based on domain knowledge and literature review [1]. The ABC model was evaluated on an enhancer perturbation dataset consisting of 109 regulatory connections inferred from experimental data [2]. The PoPS model used fine-mapped missense variants to define their ground truth set [4]. Finally, ProGeM used two different ground truth datasets: (1) 227 metabolite GWAS hits each assigned to high-confidence causal genes based on literature evidence, and (2) 562 *cis* protein quantitative trait loci (*cis*-pQTLs) data from the INTERVAL [5] study, assuming that the most likely causal gene responsible for each *cis*-pQTL signal was the gene coding for the protein [3]. As expected, the closest gene approach produced different results in the four studies: 37% recall and 47% precision in the ABC study; 55% recall and 56% precision in the L2G study; 48% recall and 46% precision in the PoPS study. In contrast, ProGeM achieved 76% precision for metabolite GWAS hits and 69% precision for *cis*-pQTLs.

Here, we used the same INTERVAL *cis*-pQTL ground truth dataset employed by ProGeM but expanded the analysis in multiple ways. First, we fine-mapped the *cis*-pQTL signals, allowing us to consider multiple conditionally distinct causal variants for each protein in the same *cis* region. Secondly, we used colocalisation instead of simple eQTL lookup to link *cis*-pQTLs to putative target genes [6]. Third, fine-mapped eQTL data from the eQTL Catalogue allowed us to perform colocalisation at the resolution of individual signals instead of genomic regions [7,8]. Finally, identifying gene-protein pairs with two or more colocalising signals allowed us to evaluate five Mendelian randomisation (MR) approaches for causal gene prioritisation. The closest gene approach (72% precision) outperformed all colocalisation methods. Combining colocalisation with Mendelian randomisation (MR) and restricting analysis to gene-protein pairs with two or more shared signals increased the precision of target gene identification from 45% to 81%, but this came with a large decrease in recall (from 46% to 7%). The reduction in recall was primarily driven by the small sample size of eQTL datasets that limited the power to detect secondary eQTL signals. Importantly, we found that *cis*-eQTL often violated one or more MR assumptions and using robust inference methods that accounted for these violations was essential to avoid false positives.

## Results

### Overview of the experimental design

To compare different colocalization methods on real-world data, we integrated *cis*-eQTL data from 131 fine-mapped tissue-specific datasets from the eQTL Catalogue (n = 65-702 individuals, Table S1) [9] with fine-mapped plasma protein QTL data from the INTERVAL cohort (n = 3,301) [5]. We used three Bayesian colocalisation methods: **coloc.abf** [6], which assumes a single causal variant per locus; **coloc.susie** [7], which supports multiple fine-mapped causal variants; and colocalization posterior probability (**CLPP**) defined at the variant level [8]. We considered a colocalization signal to be significant in a locus if the posterior probability of colocalization (PP4) was greater than 0.8 for coloc.abf or coloc.susie. For CLPP, we used the commonly used threshold of 0.1 [4,10] as the CLPP value is not directly comparable to PP4 from coloc.abf and coloc.susie.

To illustrate how the three colocalisation methods work, we looked at the colocalisation between *POSTN* gene expression in the GTEx fibroblast dataset (QTD000216, n = 483) and POSTN plasma protein abundance in the INTERVAL dataset. The colocalisation was clearly detected by coloc.abf (PP4 = 0.975) (Fig 1B). Interestingly, at this locus coloc.susie detected a colocalisation for two independent fine-mapped signal pairs. The first eQTL signal colocalised with the second pQTL signal (PP4 = 0.979) (Fig 1C) and the second eQTL signal colocalised with the fifth pQTL signal (PP4 = 0.944) (Fig 1D). The two independent signals can also be seen on the coloc.abf plot (Fig 1B), where they had proportional effects on gene and protein levels and thus did not interfere with the single causal variant assumption of coloc.abf. CLPP did not detect a colocalisation at this locus, because for the first fine-mapped signal pair, the CLPP value was below the 0.1 threshold (Fig 1C) and for the second signal pair, SuSiE did not detect a credible set for the fifth pQTL signal.

**Figure 1:**
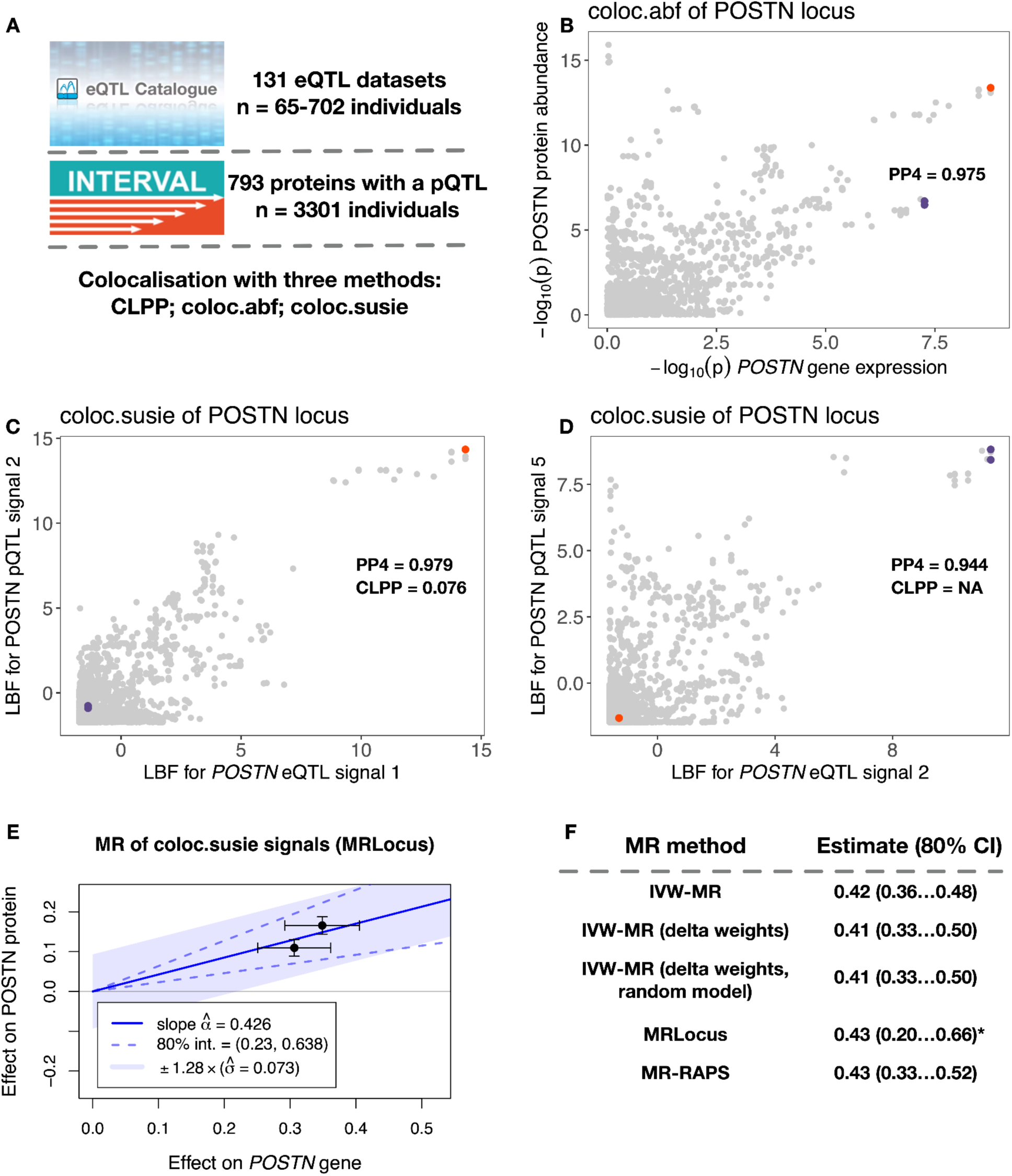
Overview of datasets and analysis methods. (**A**) Fine-mapped *cis*-eQTLs from 131 distinct datasets and *cis*-pQTLs from the INTERVAL cohort were retrieved from eQTL Catalogue release 6. The coloc.abf, coloc.susie and CLPP methods were used to identify colocalising *cis*-eQTLs and *cis*-pQTLs pairs. (**B**) coloc.abf colocalisation between *POSTN* gene expression in GTEx fibroblasts (QTD000216, n = 483) and POSTN protein abundance in plasma. (**C-D**) coloc.susie colocalisation for two pairs of fine-mapped eQTL-pQTL signals in the same datasets. The red dot represents the shared lead QTL variant for the first eQTL signal and the second pQTL signal. The two purple dots represent the strongly linked (r^2^ = 0.995) lead variants for the second eQTL signal and the fifth pQTL signal. Instead of marginal association summary statistics, coloc.susie uses log Bayes factors (LBFs) for each fine-mapped signal. (**E**) MR with the pair of colocalising eQTLs from panels C and D is consistent with a causal effect from gene expression to protein abundance (graph created by MRLocus). (**F**) All five MR methods produce similar effect estimates and confidence intervals for the example on panel E. *MRLocus provides a Bayesian credible interval instead of a confidence interval.

For the 96 gene-protein pairs for which coloc.susie detected multiple independent colocalising signals in a single dataset, we further tested five Mendelian randomization (MR) methods (Figure 1E-F) to check if the effect sizes of the distinct colocalising genetic signals were consistent with a putative causal effect of gene expression on protein abundance. We started with inverse variance weighted Mendelian randomisation (IVW-MR) implemented in the MendelianRandomization R package [11] and the IVW-MR with delta weights [12] to account for the uncertainty of instrument effects on exposure. To account for potential violations of MR assumptions in the eQTL data, we also included three other methods that include additional modelling considerations: multiplicative random-effects IVW-MR that models overdispersion heterogeneity between instruments [12,13], MRLocus that models the dispersion of instrument’s effects via the allelic spread parameter [14], and MR-RAPS that models overdispersion heterogeneity while also accounting for outlier instruments [15]. In the case of POSTN, all MR methods yielded very similar causal effect estimates and confidence intervals (Fig 1F).

### Evaluating colocalisation for causal gene identification

We first counted the number of proteins that were found to colocalize with any gene in any of the eQTL datasets. Since coloc.abf is only able to work with the strongest signal in a locus, we restricted coloc.susie to use only the first pQTL signal and CLPP to use only the first pQTL credible set for each protein for an objective comparison of the three methods. Even with these restrictions, coloc.susie found the largest number of proteins to colocalize (482 of the 793 tested), 57 of which were not discovered using the other two methods (Fig 2A). CLPP, on the other hand, found just 183 colocalizations, all of which were also detected by the other methods. Considering all independent pQTL signals increased the advantage of coloc.susie even further (Fig S1).

**Figure 2:**
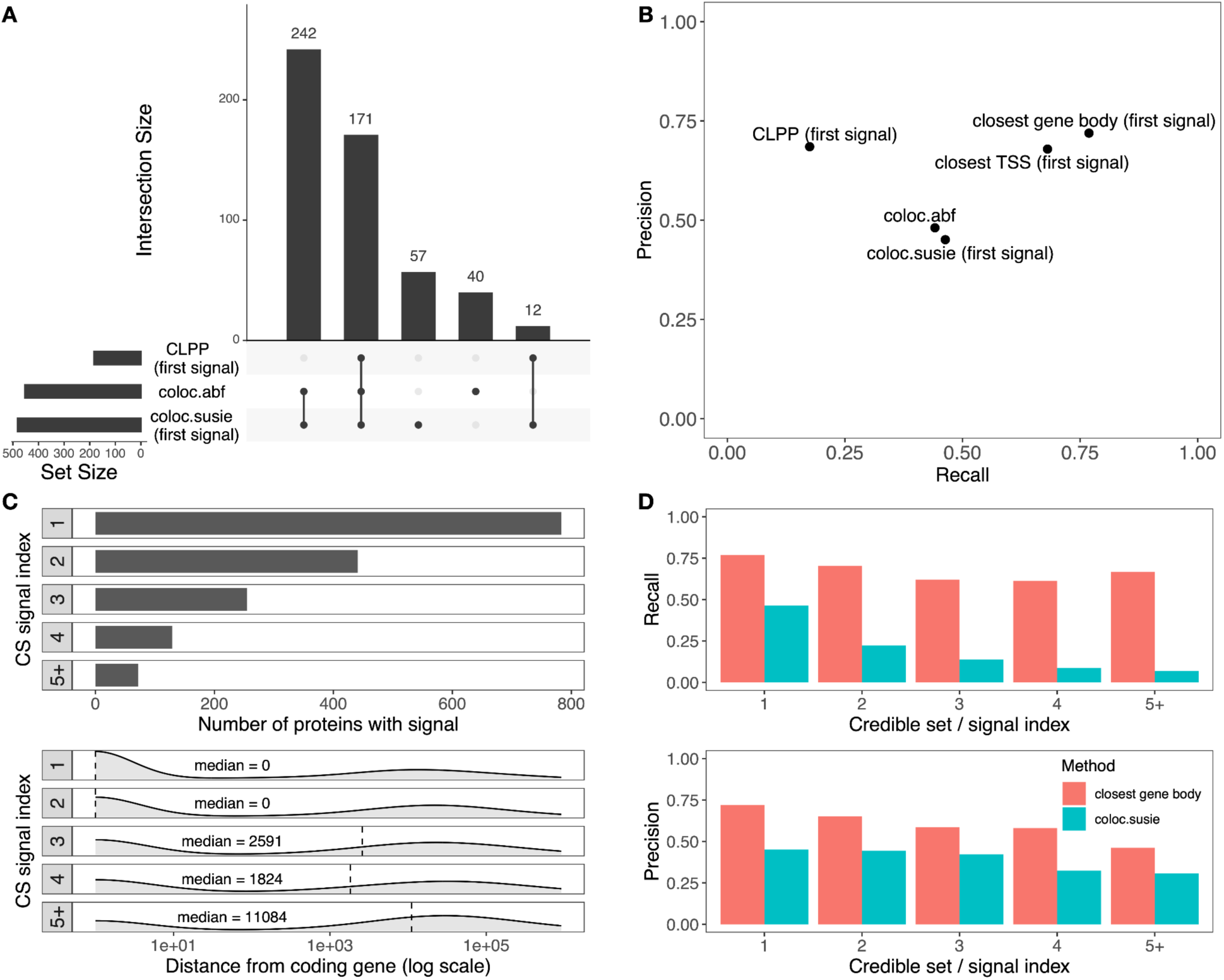
Comparing the performance of three colocalization methods in causal gene identification. (**A**) The histogram on the left shows the number of proteins with at least one colocalising eQTL detected by the CLPP, coloc.abf and coloc.susie methods. The histogram on the right is the UpSetR [16] plot showing the overlap of the colocalization events detected by the three methods. For CLPP and coloc.susie, only the first fine-mapped pQTL signal was included in the analysis. (**B**) The precision and recall of the three colocalization methods in causal gene identification relative to the closest gene approach. (**C**) Overview of the independent fine-mapped pQTL credible sets detected for each protein. The top histogram shows the number of proteins with 1,2,3,4,5 or more credible sets detected. The density plots at the bottom show the distance from the fine-mapped pQTL credible set lead variant to the gene body of the corresponding protein coding gene. (**D**) The precision and recall of the closest gene and coloc.susie methods as a function of the credible set index. CLPP - colocalization posterior probability.

Relying on the central dogma, we considered only significant colocalization signals between a *cis*-pQTL and the gene coding for the protein to be true positives (TP) and all other significant signals to be false positives (FP). If a protein had at least one significant pQTL in the dataset but a colocalizing signal between it and its coding gene was not found, this was considered a false negative (FN). This allowed us to assess the recall (TP/(TP+FN)), the percentage of analysed proteins found to colocalize with the gene coding for them) and the precision (TP/(TP+FP)), the percentage of correct protein-gene pairs among all unique protein-gene pairs) of each method.

Using only the first signal for each protein, coloc.susie found the correct gene for 367 of the 793 proteins with purity filtered credible sets (46.3% recall). Coloc.abf was able to match a similar percentage of them to the coding gene (44.1%) at a slightly higher precision (48.1% vs 45.1%) (Fig 2B). The variant-based approach of CLPP was the most precise of the three (68.5%), but yielded the correct gene for less than a fifth of the proteins (17.5%) (Fig 2B). However, all three colocalization methods were outperformed by a simple heuristic that assigned each pQTL to the gene body of the closest protein-coding gene (76.9% recall, 71.9% precision) (Fig 2B). Notably, using distance to the closest transcription start site (TSS) instead of gene body decreased both precision (67.9%) and recall (68.1%), suggesting that some pQTLs might alter protein abundance in a TSS-independent manner (e.g. missense or 3′ UTR variants altering protein or mRNA stability).

We then speculated that the good performance of the closest gene approach could be caused by the strongest pQTLs being located close to or within their target genes. Indeed, the median distance from the first fine-mapped pQTL signal to the corresponding protein coding gene was 0 bp, meaning that most primary pQTLs were located within the gene body of the corresponding gene. This increased to 11,084 bp for the fifth and further signals with a wide spread (Fig 2C). We observed that both the precision (71.9% vs 58.5%) and recall (76.9% vs 62.0%) of the closest gene approach decreased slightly for tertiary and further pQTL signals (Fig 2D). The precision of the coloc.susie method was less affected by the pQTL signal index, but still always remained below the closest gene approach (e.g. 42.2% vs 58.5% for third signals). Furthermore, recall of the colocalization approach decreased significantly for secondary pQTL signals (Fig 2D), suggesting that there might be less power to detect colocalisations at secondary signals due to their smaller effect sizes.

### Evidence from multiple colocalising eQTLs improves precision

Fine-mapping allowed us to identify multiple conditionally distinct *cis*-QTLs for both proteins and genes. In the INTERVAL dataset, we detect two or more pQTLs for 445 (56%) proteins with 71 (9%) having five or more independent signals in the *cis* region (Fig 3A). In the eQTL Catalogue, the number of genes with multiple independent eQTLs depended on the sample size, but even in the group of datasets with the largest sample size (n > 350) only 20% of the genes had multiple independent fine-mapped eQTLs (Fig 3B). This suggests that most current eQTL datasets are too small to effectively fine-map multiple independent signals. This is consistent with previous analyses conducted by the GTEx, MetaBrain and AdipoExpress projects, where the number of secondary eQTLs was strongly dependent on the eQTL sample sizes and reached more than 50% for tissues with the largest sample size [17–19].

**Figure 3:**
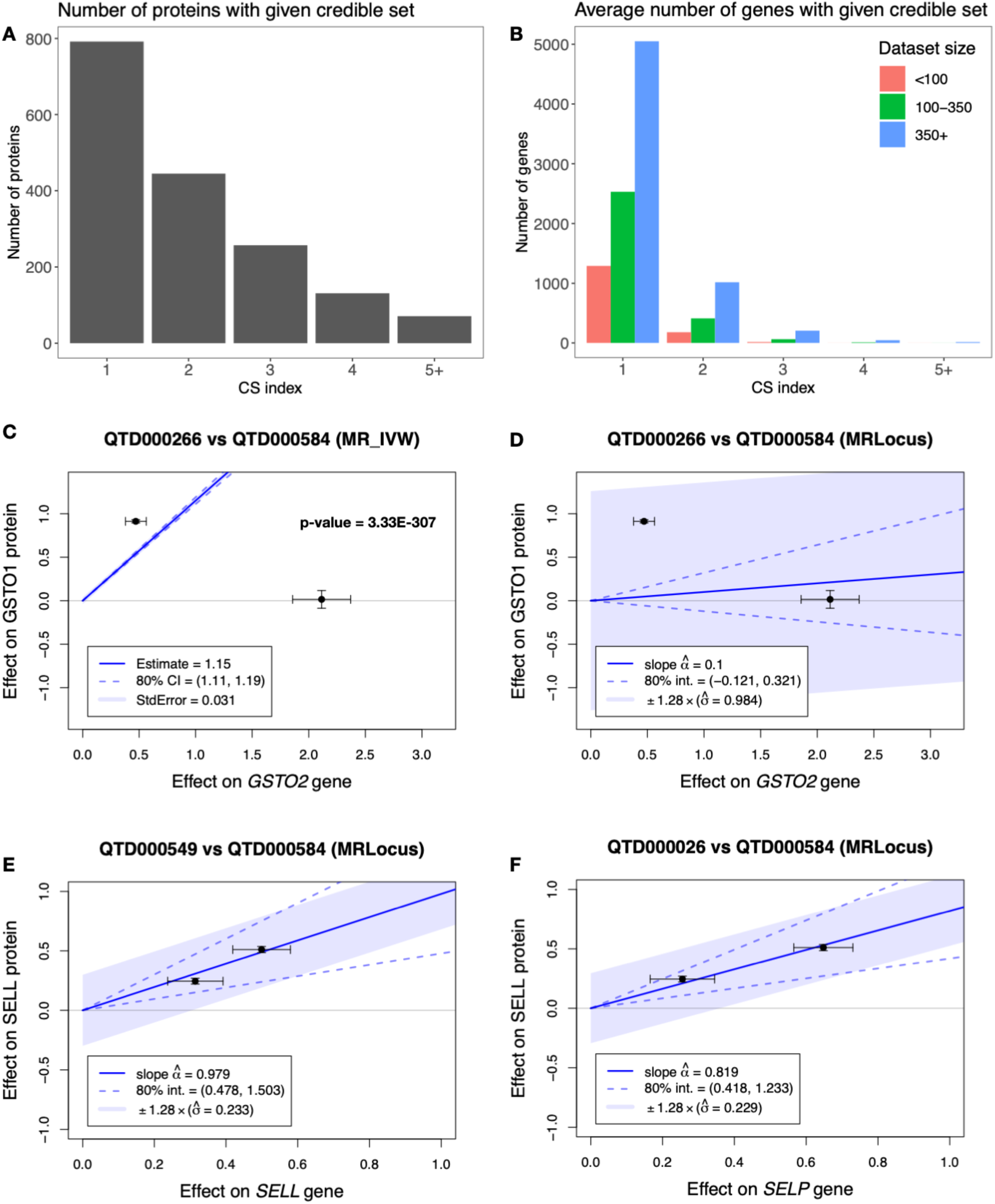
Using Mendelian randomisation to assess effect size concordance between colocalising signal pairs. (**A**) Number of proteins with evidence of multiple signals (as indicated by credible set (CS) index) in the INTERVAL study (n = 3,301). (**B**) Average number of genes with evidence of multiple signals (as indicated by CS index), stratified by the eQTL Catalogue dataset sample size. (**C**) Two colocalising QTL signals between *GSTO2* gene expression in the liver (QTD000266, n = 208) and GSTO1 protein abundance in plasma. Standard IVW-MR detects a highly significant effect (p-value < 3*10^-290^) that is not supported by the data. (**D**) The signal from panel C analysed with MRLocus. MRLocus 80% credible interval overlaps zero with wide allelic spread, because the two QTLs have inconsistent effects on gene expression and protein abundance. (**E**) Significant MR signal between *SELL* gene expression in blood (QTD000549, n = 195) and SELL protein abundance in plasma. (**F**) Significant MR signal between *SELP* gene expression in BLUEPRINT neutrophils (QTD000026, n = 196) and SELL protein abundance in plasma.

Coloc.susie is able to consider all pairwise colocalisations between independent fine-mapped QTLs in the same *cis* region (Fig 1C-D). Across all eQTL Catalogue datasets, we detected 321 gene-protein pairs with two or more colocalising QTLs in the same *cis* region. The same gene-protein pair was often detected in multiple eQTL datasets, with 96 of them being unique (Table 1). These low numbers are primarily caused by a lack of secondary fine-mapped eQTL signals detected in the eQTL Catalogue (Fig 3B). As a result, this approach had a recall of only 8.6%, but precision increased from 45.1% (Fig 2B) to 70.8% (Table 1), suggesting that observing multiple colocalising signals can significantly improve the precision of target gene identification.

**Table 1:** Comparison of the MR methods used. TP with positive slope - the fraction of true positive gene-protein-dataset triplets for which MR fitted a positive slope. FP with positive slope - the fraction of false-positive gene-protein-dataset triplets for which MR fitted a positive slope. While precision and recall are calculated at the level of gene-protein pairs, the direction of the MR slope is estimated separately in each gene-protein-dataset triplet.

| Method | Precision | Recall | TP with positive slope | FP with positive slope |
| --- | --- | --- | --- | --- |
| coloc.susie (multiple signals), no MR | 68/96 (70.8%) | 68/793 (8.6%) | NA | NA |
| IVW-MR | 65/92 (70.7%) | 65/793 (8.2%) | 240/263 (91.3%) | 25/48 (52.1%) |
| IVW-MR (delta weights) | 63/88 (71.6%) | 63/793 (7.9%) | 232/252 (92.1%) | 23/44 (47.7%) |
| IVW-MR (delta weights, random model) | 56/69 (**81.1%**) | 56/793 (7.1%) | 205/215 (95.3%) | 10/24 (41.7%) |
| MRLocus | 48/63 (76.2%) | 48/793 (6.1%) | 196/203 (**96.6%**) | 10/21 (47.6%) |
| MR-RAPS | 64/87 (73.6%) | 64/793 (8.1%) | 229/254 (90.2%) | 22/43 (51.2%) |

### Effect size concordance between multiple colocalising QTLs

When focussing on gene-protein pairs with two or more independent colocalising QTL signals, we noticed that the effect sizes of these QTLs were often discordant (Fig 3C, Fig S2). We hypothesised that excluding these strongly discordant gene-protein pairs that are incompatible with a causal effect from gene expression to protein abundance might further increase precision. We first used inverse-variance weighted Mendelian randomisation (IVW-MR) to identify these discordant pairs. Unexpectedly, we found that filtering colocalization results for significant MR slope (80% confidence interval not intersecting zero) had only minimal effect on identified gene-protein pairs (92 of 96 remained) and virtually no effect on precision and recall (Fig 3C, Table 1). These results remained robust to using more stringent filtering criteria (e.g. all 92 pairs remained at FDR < 1%). Using IVW-MR with delta weights to better model large standard errors on the exposure typical for eQTL data also had minimal effect (Table 1). When consulting the literature, we realised that this behaviour is likely driven by an assumption of the standard IVW-MR model that all instruments included in the analysis provide a (noisy) estimate of the same underlying causal effect. Violations of this assumption (Fig 3C, Fig S2) can lead to underestimation of standard errors (Fig 3C) and overestimation of statistical significance [12,15,20].

We tested three robust MR methods that explicitly model overdispersion heterogeneity or dispersion of instrument’s effects: multiplicative random-effects IVW-MR [12], MRLocus [14] and MR-RAPS [15]. For the *GSTO2*-GSTO1 example (Fig 3C), both the random-effects IVW-MR and MRLocus now correctly inferred a null effect while MR-RAPS still detected a significant effect (Fig 3D, Fig S3). The same was true for several other examples (Fig S4). Overall, random-effect IVW-MR (81.1% precision, 7.1% recall) performed slightly better than MRLocus (76.2% precision, 6.1% recall) while MR-RAPS (73.6% precision, 8,1% recall) performed similarly to the standard IVW-MR method (Table 1). On closer inspection, the poor performance of MR-RAPS seemed to be caused by its outlier detection feature that often excluded one of the two instruments from the analysis (Fig S3-4).

### Concordance in gene and protein effect size direction

An assumption that we can make when working with gene-protein pairs is that for true causal relationships, the variants that increase gene expression should also increase protein abundance (i.e. have a positive MR slope). We split the gene-protein pairs identified by the five MR methods into true positives if the gene coded for the protein and false positives otherwise. We found that MRLocus had the highest fraction of true positive pairs with a positive slope (96.6%) closely followed by random-effect IVW-MR (95.3%) (Table 1). The other three methods had lower effect size direction concordance that ranged from 90.2% (MR-RAPS) to 92.1% (delta-weighted IVW-MR) (Table 1), indicating that explicit modelling of overdispersion heterogeneity not only increases precision, but the gene-protein pairs detected with robust methods (MRLocus and random-effect IVW-MR) also more often display the expected direction of effect. In contrast, ∼50% of the false positive gene-protein pairs from the five MR methods had a positive slope. In some cases, the remaining false positives from the MRLocus and random-effect IVW-MR analysis with positive slopes reflected neighbouring genes where the same instruments had highly concordant effects on both genes, highlighting how strong local co-regulation can confuse even the best causal inference approaches (Figure 3E-F).

### Increasing the power of *cis*-MR via eQTL meta-analysis

Although MR with two or more colocalising signals increased the precision of target gene identification to 81%, this came at the cost of significantly reduced recall (7.1%). The primary reason for detecting few multi-signal gene-protein pairs in our analysis is the relatively small sample size of eQTL datasets (n = 65-702) in the eQTL Catalogue. To test if the recall of the MR methods could be increased by meta-analysing eQTLs across multiple studies from the same tissue, we obtained the *cis*-eQTL summary statistics from the AdipoExpress project, a meta-analysis of five subcutaneous adipose tissue studies (n = 2,344). Although AdipoExpress was not able to use SuSiE for fine-mapping due to the risk of false positives [21], they used all-but-one conditional analysis [22] to identify conditionally distinct signals. After converting these conditional summary statistics to approximate Bayes factors (see Methods), we performed colocalisation with INTERVAL pQTLs using the same workflow that we previously used for the eQTL Catalogue datasets.

We compared the colocalisation that we detected in the AdipoExpress dataset with those from the best-powered adipose tissue dataset from the eQTL Catalogue (TwinsUK, n = 381). We found that the number of colocalising gene-protein pairs increased by approximately 2-fold for both coloc.abf (from 98 to 195) and coloc.susie (104 to 242, first signal only). Consistent with the observation that larger sample sizes increase the power to detect secondary eQTL signals (Fig 3B) [17–19], the number of multi-signal gene-protein pairs increased from 8 to 35 (4.4-fold), but only 1 multi-signal pair was shared between the two analyses. Furthermore, 18/35 gene-protein pairs also had a significant MR effect in the AdipoExpress dataset (95% precision) (Table S2) and 7/8 had a significant MR effect in the TwinsUK dataset (87% precision) (Table S3). However, overall recall remained low (2.3% for AdipoExpress and 0.9% for TwinsUK, Table S2-3), potentially because adipose tissue is unlikely to be the causal tissue for many plasma *cis*-pQTLs.

## Discussion

A fundamental problem in human complex traits genetics is linking primarily non-coding GWAS hits to their causal target genes. Here, we used fine-mapped *cis*-pQTLs to systematically evaluate the performance of eQTL colocalisation methods in identifying causal target genes. Our key assumption was that the causal gene responsible for a *cis*-pQTL signal should be the gene coding for the protein. Our results indicate that eQTL colocalisation approaches, when performed systematically against very large eQTL databases such as the eQTL Catalogue, have generally low precision (∼50%) in identifying the correct target genes. This seems to be primarily driven by horizontal pleiotropy whereby the same eQTL variants are associated with the expression level of multiple neighbouring genes. We also found that precision can be improved (up to 81%) when combining multiple colocalising QTL signals in a Mendelian randomization framework to explicitly consider the concordance of the causal effect estimates provided by independent genetic variants. This agrees with other recent studies, affirming that combining colocalisation with Mendelian randomisation reduces confounding by LD and improves the sensitivity and specificity of identifying biologically relevant targets [23–25].

In our analysis, the closest gene approach had very high precision (71.9%), which is higher than typically seen in other studies that benchmark methods for causal gene prioritisation in the GWAS setting (range 46%-56%) [1,2,4]. Part of the reason could be that primary pQTLs might be much closer to their target genes than typical GWAS hits are (Figure 1C), matching a similar observation for eQTL variants and GWAS hits [26]. Indeed, we observed that the precision of the closest gene approach dropped to 58.5% for tertiary and further pQTL signals that were more often located outside the gene body (Fig 1D). Secondly, our closest gene approach was based on the distance to the gene body as opposed to the closest transcription start site (TSS) chosen by some other studies [1]. Indeed, using TSS instead of gene body to define the closest genes decreased the precision to 67.8% (Fig 2B).

Our results highlight the challenges of using gene expression levels as exposures in Mendelian randomisation. The primary concern is that eQTL variants often do not satisfy the exclusion restriction assumption of the MR framework [27], which states that genetic variants affect the outcome only through their effect on the gene expression level (exposure) included in the model. This assumption can be violated in at least two ways. First, we might be looking at the right gene but in the wrong context. The true causal effect of the gene expression on the outcome might be mediated in some other tissue, cell type or context that was not included in the analysis. In this scenario, failure to model *allelic spread* or *overdispersion heterogeneity* may inflate the significance of the standard IVW-MR estimates [13,14]. Secondly, due to horizontal pleiotropy, we might be looking at the wrong gene and the actual causal effect might be mediated by another gene for which the eQTL variants have highly correlated effects (e.g. *SELL* and *SELP* genes on Fig 3E-F). Thus, we caution against interpreting significant *cis*-MR slopes as direct evidence of the causal effect of gene expression levels on the outcome. Rather, we prefer to use MR to exclude exposures that are clearly *inconsistent* with a causal effect in the tested cell type or tissue. Finally, our results reinforce the need to include both positive and negative controls in MR analysis and use visualisation approaches to assess model fit [28].

A promising approach that we did not evaluate here is multivariable MR which jointly models the expression levels of all neighbouring genes [13,29]. However, multivariable MR requires that the number of genetic instruments included in the analysis equals or exceeds the number of exposures, which is unrealistic for *cis*-eQTLs from large compendia containing hundreds of cell types and tissues [30]. Furthermore, multivariable MR can identify the correct causal gene only if the right gene in the right context (cell type or tissues), or a sufficient proxy context, is included in the model as one of the exposures. There also needs to be sufficient *phenotypic heterogeneity* between the different exposures included in the model (i.e. genetic variant effects vary between the different genes) [13]. Thus, multivariable MR is unlikely to completely resolve the exclusion restriction assumption violations that we have observed here.

Choosing the closest gene almost always outperformed eQTL colocalisation when identifying causal genes responsible for *cis*-pQTLs. Furthermore, even though MR with multiple independent eQTLs did outperform the closest gene approach in terms of precision (81.2% vs 71.9%), this came at the cost of a significant reduction in recall (76.9% vs 7.1%). This reduction in recall was primarily driven by limited power to detect secondary eQTL signals in existing datasets (Fig 3B). Consistent with this hypothesis, we found that using adipose tissue *cis*-eQTL conditional meta-analysis summary statistics from the AdipoExpress project (n=2,344) instead of TwinsUK (n = 415) increased the recall of eQTL *cis*-MR by 2.5-fold from 0.9% to 2.3% (Table S2-3). Thus, a promising avenue to improve the recall of eQTL MR is to increase the sample sizes of eQTL datasets by either collecting new samples or performing meta-analysis across multiple existing datasets. A potential added benefit is that secondary eQTLs might represent more distal context-specific effects that are more likely to overlap disease GWAS hits [26]. Finally, successful target gene prioritisation will likely require triangulation of evidence from multiple genetic and non-genetic sources. Fortunately, multiple competing statistical models are currently actively being developed to support this integration (e.g. L2G [1] and PoPS [4]).

## Methods

### Datasets used in the analysis

We downloaded eQTL summary statistics and fine mapping results for 34 studies [18,31–59] from the eQTL Catalogue (release 6) FTP server (https://www.ebi.ac.uk/eqtl/Data_access/). The genotype and protein abundance data from the INTERVAL cohort [5] were downloaded from EGA (accessions EGAD00010001544 and EGAD00001004080) after access was approved by the ‘Plasma pQTLs in INTERVAL cohort’ data access committee.

### INTERVAL pQTL data processing

#### Genotype imputation

The imputed genotypes from the INTERVAL cohort were based on the GRCh37 coordinates, but all fine mapping results from the eQTL Catalogue used GRCh38 coordinates. We initially used CrossMap.py [60] to convert imputed genotypes to GRCh38 coordinates, but found that this approach caused some artefactual fine-mapping results due to variants lost in the lift-over process. To avoid these issues, we extracted genotyped variant positions of Affymetrix Axiom UK Biobank array from the INTERVAL imputed genotype files and re-imputed genotypes to the 1000 Genomes 30x on GRCh38 reference panel with the <u>eQTL-</u> <u>Catalogue/genimpute</u> v23.07.1 workflow. The same workflow was previously used to impute genotypes for all eQTL Catalogue datasets that used genotyping microarrays [9].

#### Protein data processing and association testing

We downloaded the pre-processed SomaLogic protein abundance data and aptamer metadata from EGA (EGAD00001004080). We applied inverse normal transformation to the protein abundance data and used the g:Profiler [61] web tool to map SomaLogic protein names to Ensembl gene ids. The metadata for SomaLogic aptamers and their mapping to Ensembl gene ids can be downloaded from Zenodo (https://doi.org/10.5281/zenodo.7808390). The *cis*-pQTL analysis and fine mapping were conducted using the <u>eQTL-Catalogue/qtlmap</u> v23.02.1 workflow as described previously [9]. In all downstream analyses, we used 793 proteins that had at least one purity-filtered SuSiE credible set.

### AdipoExpress data processing

The AdipoExpress project performed *cis*-eQTL meta-analysis across five subcutaneous adipose tissue studies (total n = 2,344) [19]. We downloaded AdipoExpress summary statistics from https://mohlke.web.unc.edu/data/adipoexpress/. Since AdipoExpess analysis used the GRCh37 reference genome, we first converted the variant positions to GRCh38 coordinates with the MungeSummstats [62] R package. Since using SuSiE to identify conditionally distinct signals is prone to false positives in a meta-analysis setting [21], AdipoExpress used all-but-one conditional meta-analysis and also released summary statistics for distinct signals conditioned on all other significant signals in the same *cis-*region. To use these results with coloc.susie, we first converted the all-but-one conditional betas and standard errors to log approximate Bayes factors (LABFs) using the process.dataset() function from the coloc R package. Subsequently, we used the LABFs in place of the log Bayes factors in the coloc.susie method.

### Colocalization between *cis*-eQTLs and *cis*-pQTLs

We downloaded fine-mapped *cis*-eQTL summary statistics for 131 datasets of the eQTL Catalogue release 6 [9] from eQTL Catalogue FTP server. We ran colocalization analyses pairwise between all eQTL datasets and pQTL data from the INTERVAL study. We set the *cis*-window for each locus at 2 million base pairs centred at the transcription start site. The Nextflow workflow implementing the CLPP, coloc.susie and coloc.abf colocalisation methods is available from GitHub (https://github.com/ralf-tambets/coloc). The workflow assigned colocalization probabilities for each *cis-*pQTL locus in the INTERVAL dataset and each *cis*-eQTL locus of each eQTL dataset. We only included protein coding genes in the analysis as these are much more likely to be the causal genes for pQTLs. We also excluded all protein complexes from the INTERVAL dataset as their abundance could be influenced by all of their constituents independently [63,64].

#### CLPP

We calculated CLPP as described previously [10]. Briefly, we joined the data from the eQTL study and the pQTL study by the variant name. We calculated CLPP for each variant by multiplying the posterior inclusion probabilities from both studies, and summed the resulting values up for each credible set in the eQTL dataset. A signal was significant if CLPP exceeded 0.1. This approach yielded 2,278 unique colocalizing gene-protein-dataset triplets (Table S2).

#### coloc.abf

We analysed each dataset chromosome by chromosome by running coloc.abf [6] on summary statistics (beta, standard error, MAF) between each eQTL gene and a subset of the *cis*-pQTLs that fell in the *cis*-window, unless more than 90% of the variants in the pQTL gene fell outside the *cis*-window. The prior probabilities that a SNP is associated with either trait were set at 1×10^-4^ and the prior probability that a SNP is associated with both traits was set at 5×10^-6^. A signal was significant if PP4 exceeded 0.8. This approach yielded 4,890 unique colocalising gene-protein-dataset triplets (Table S3).

#### coloc.susie

Data preparation for coloc.susie [7] was similar to that of coloc.abf with the exception that the input data consisted of SuSiE log Bayes factors (LBFs) for all fine-mapped signals instead of marginal betas and standard errors. We used the coloc.bf_bf function to calculate the colocalisation posterior probabilities, which we ran with the same prior probabilities of association as for coloc.abf. A signal was significant if PP4 exceeded 0.8. This approach yielded 8,501 unique colocalizing gene-protein-dataset triplets (Table S4).

### Summary analysis

To determine the closest gene to a given protein for benchmarking purposes, we found the lead variant for each credible set based on z-score and calculated the distance from it to the start and end coordinates of each protein coding gene. If the lead variant fell within the gene body, the distance to the gene was set to zero. In cases of equal closest distances, all tied genes were considered closest, with at most one of them being a true positive.

### Mendelian randomization

We tested five different MR methods: default mr_ivw() method from the MendelianRandomization R package version 0.9.0 [11]; the same function with the weights argument set to “delta” [12]; the same function with the weights argument set to “delta” and the model argument set to “random”; the mr_raps() function from the MR-RAPS R package version 0.4.1 [15] with the over.dispersion argument set to TRUE and the loss.function argument set to “tukey”; and the fitSlope() function from the MRLocus R package version 0.0.26 [14]. We considered the signal from a gene-protein-dataset triplet as significant if the 80% confidence interval did not include 0 (using MendelianRandomization and MR-RAPS) or if the 80% credible interval did not include 0 (using MRLocus).

## Data availability

All eQTL summary statistics, fine mapping results and log bayes factors are available from the eQTL Catalogue FTP server (https://www.ebi.ac.uk/eqtl/). The pQTL summary statistics and fine mapping results from the INTERVAL cohort have also been deposited to the eQTL Catalogue under the accession QTD000584. The individual-level genotypes and protein abundances from the INTERVAL cohort were downloaded from EGA (accessions EGAD00001004080 and EGAD00010001544).

## Author contributions

R.T. performed all colocalisation and Mendelian randomisation analyses presented in the paper. A.K. prepared the INTERVAL proteomics dataset for pQTL analysis. P.K. developed the genotype imputation workflow for low-coverage whole genome sequencing data. K.A. and R.T. wrote the manuscript with input from all authors.

## Acknowledgements

We thank S. Kasela for her helpful comments on the manuscript. The colocalisation and Mendelian randomisation analyses were performed at the High Performance Computing Center, University of Tartu.

## Supplementary Figures

**Figure S1.**
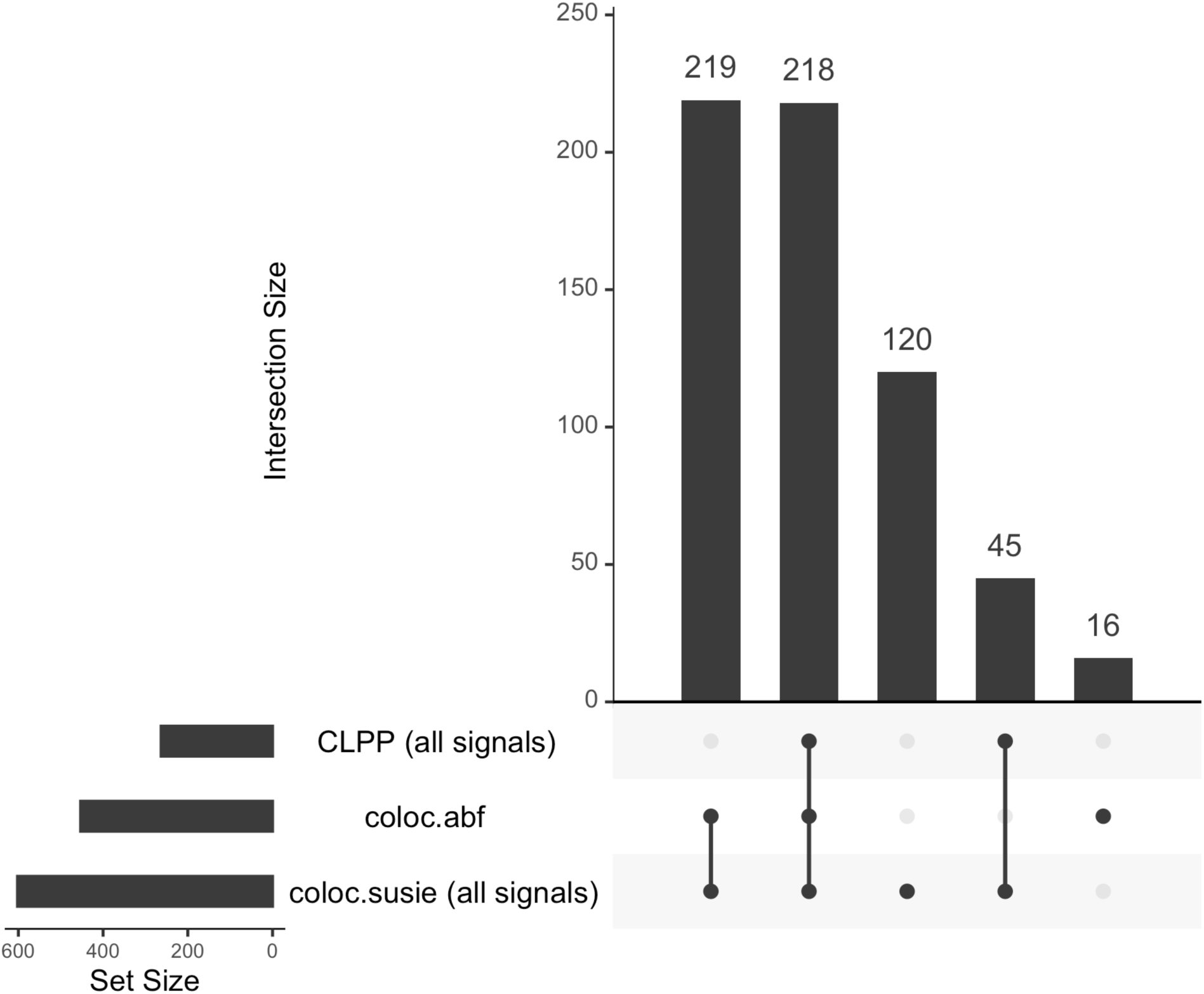
The histogram on the left shows the number of proteins with at least one colocalising eQTL detected by the CLPP, coloc.abf and coloc.susie methods. The histogram on the right is the UpSetR [16] plot showing the overlap of the colocalization events detected by the three methods. For CLPP and coloc.susie, all fine-mapped pQTL signals were included in the analysis. The total number of proteins included in the analysis was 793.

**Figure S2.**
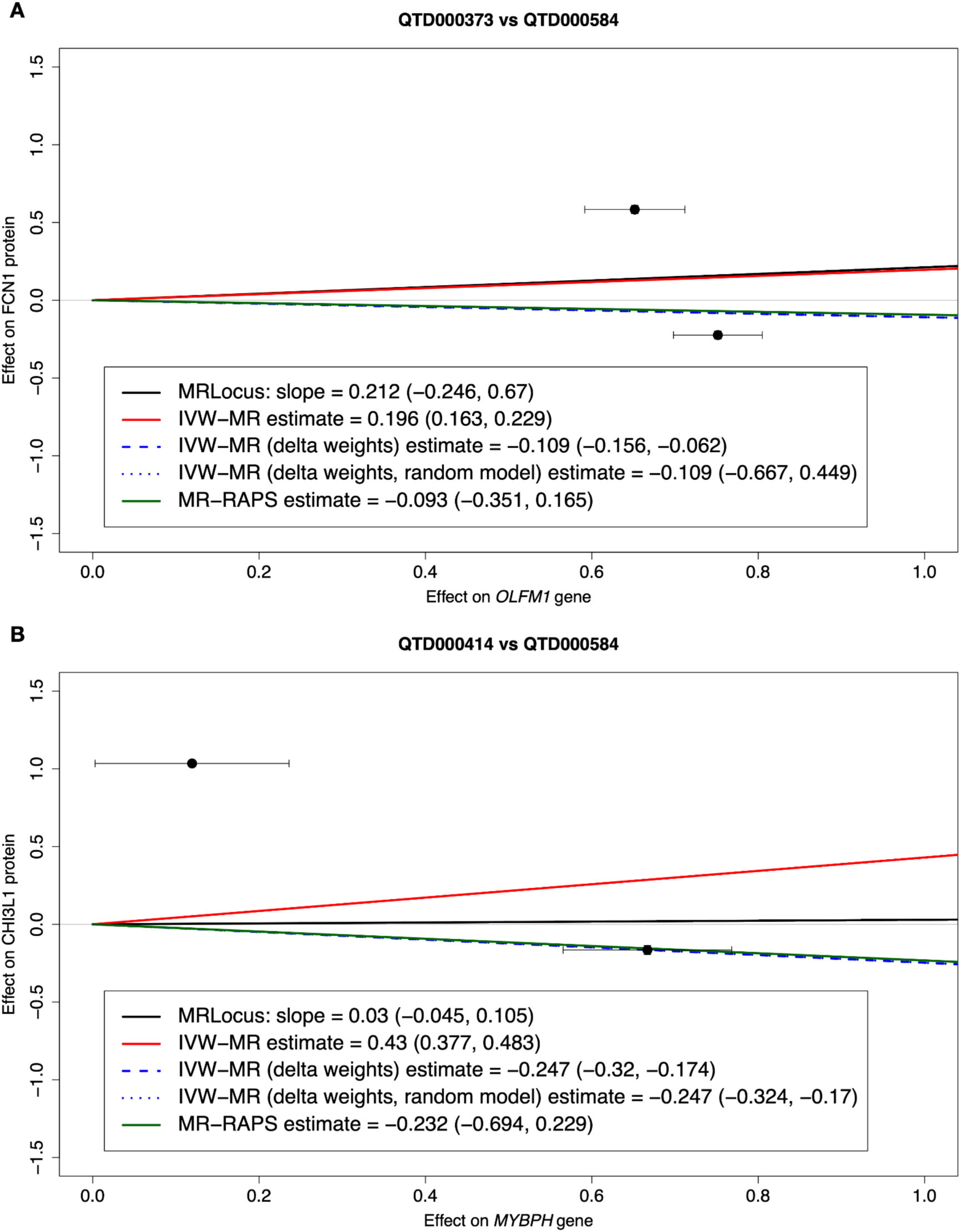
Examples of colocalising eQTLs with discordant effect sizes on protein abundance. (**A**) Discordant effect sizes between *OLFM1* gene expression in blood (QTD000373, n = 471) and FCN1 protein abundance in plasma. Bottom panel shows the MR effect size estimates and 80% confidence intervals (80% credible interval for MRLocus) for all five MR methods. (**B**) Discordant QTL effect sizes between *MYBPH* gene expression in LPS-stimulated monocytes (QTD000414, n = 184) and CHI3L1 protein abundance in plasma. Bottom panel shows the MR effect size estimates and 80% confidence intervals (80% credible interval for MRLocus) for all five MR methods.

**Figure S3.**
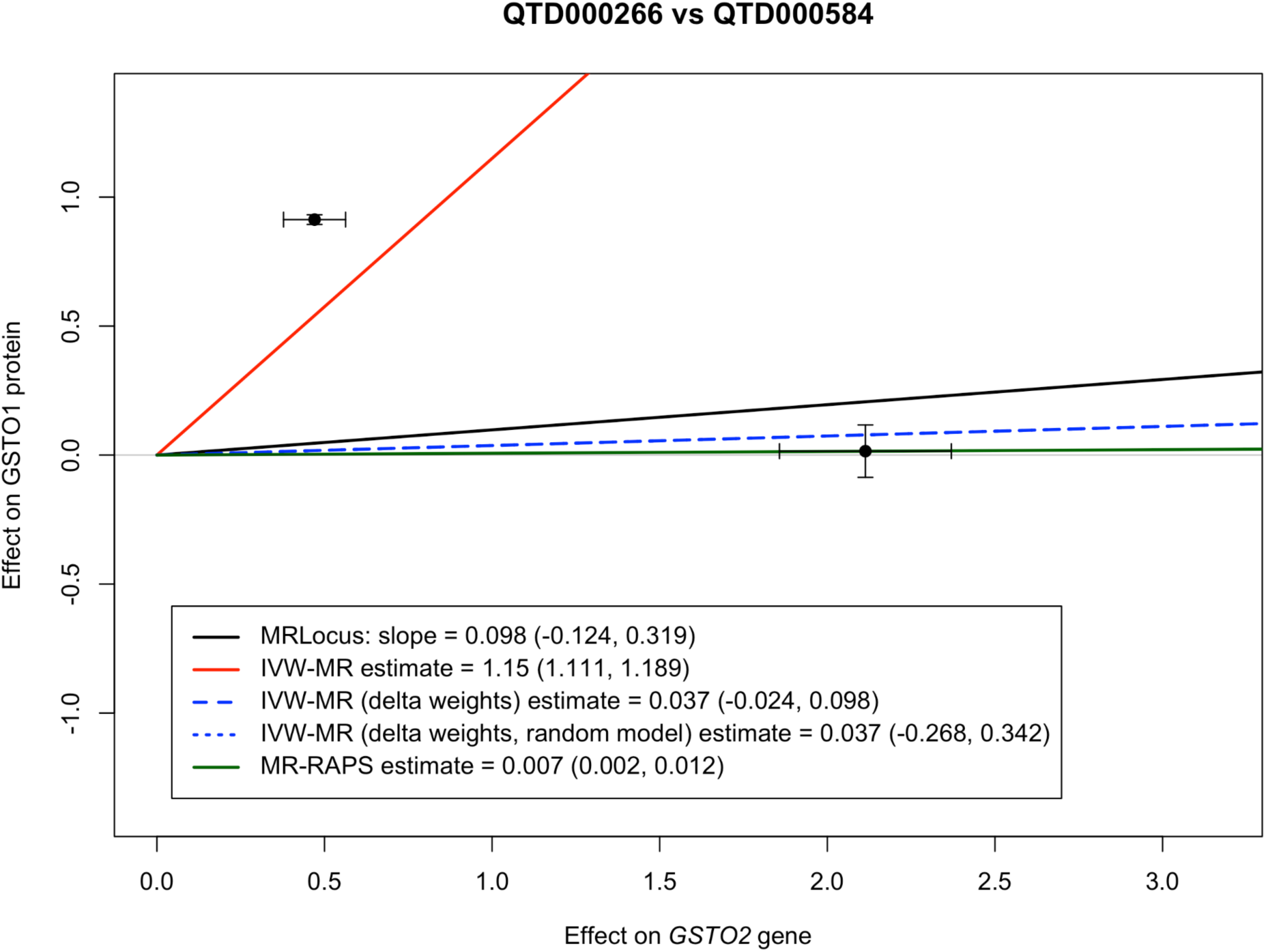
Mendelian randomisation between *GSTO2* gene expression in liver (QTD000266) and GSTO1 protein abundance in plasma using the five MR methods included in the study. IVW-MR infers a very strong positive effect. IVW-MR with delta weights detects a null effect with a narrow 80% confidence interval. Switching to random-effect IVW-MR gives the same causal effect estimate but a much larger confidence interval. This behaviour is also expected from theory [12]. MRLocus gives a similar estimate and 80% credible interval to the random-effect IVW-MR model. MR-RAPS excludes the instrument with a large effect on protein abundance as an outlier and infers a small but very precise effect based on the other instrument alone.

**Figure S4.**
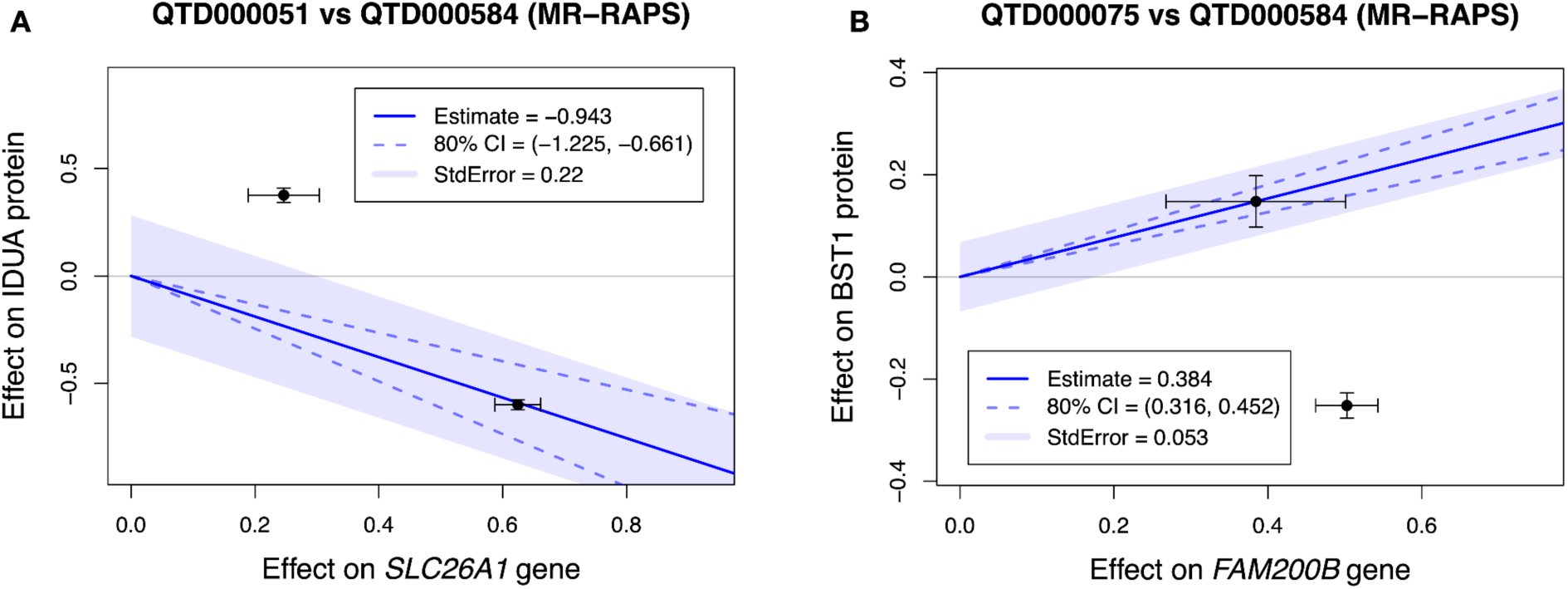
Additional examples of colocalising eQTLs with discordant effect sizes on protein abundance, analysed using MR-RAPS. (**A**) Mendelian randomisation between *SLC26A1* gene expression in brain tissue (QTD000051, n = 479) and IDUA protein abundance in plasma. (**B**) Mendelian randomisation between *FAM200B* gene expression in ileum tissue (QTD000075, n = 586) and BST1 protein abundance in plasma. In both cases, MR-RAPS discards one variant as an outlier and bases its estimations on the other.

**Table S2:** Comparison of the MR methods on AdipoExpress dataset. TP with positive slope - the fraction of true positive gene-protein-dataset triplets for which MR fitted a positive slope. FP with positive slope - the fraction of false-positive gene-protein-dataset triplets for which MR fitted a positive slope.

| Method | Precision | Recall | TP with positive slope | FP with positive slope |
| --- | --- | --- | --- | --- |
| coloc.susie (multiple signals), no MR | 29/35 (76.3%) | 29/793 (3.7%) | NA | NA |
| IVW-MR | 29/35 (82.9%) | 29/793 (3.7%) | 28/29 (96.6%) | 2/6 (33.3%) |
| IVW-MR (delta weights) | 29/35 (82.9%) | 29/793 (3.7%) | 28/29 (96.6%) | 2/6 (33.3%) |
| IVW-MR (delta weights, random model) | 18/19 (**94.7%**) | 18/793 (2.3%) | 17/18 (94.4%) | 0/1 (0.0%) |
| MR Locus | 20/23 (87.0%) | 20/793 (2.5%) | 19/20 (95.0%) | 1/3 (33.3%) |
| MR-RAPS | 27/31 (87.1%) | 27/793 (3.4%) | 26/27 (96.3%) | 1/4 (25.0%) |

**Table S3:** Comparison of the MR methods on TwinsUK adipose dataset. TP with positive slope - the fraction of true positive gene-protein-dataset triplets for which MR fitted a positive slope. FP with positive slope - the fraction of false-positive gene-protein-dataset triplets for which MR fitted a positive slope.

| Method | Precision | Recall | TP with positive slope | FP with positive slope |
| --- | --- | --- | --- | --- |
| coloc.susie (multiple signals), no MR | 7/8 (87.5%) | 7/793 (0.9%) | NA | NA |
| IVW-MR | 7/8 (87.5%) | 7/793 (0.9%) | 7/7 (100%) | 1/1 (100%) |
| IVW-MR (delta weights) | 7/8 (87.5%) | 7/793 (0.9%) | 6/7 (85.7%) | 1/1 (100%) |
| IVW-MR (delta weights, random model) | 7/8 (87.5%) | 7/793 (0.9%) | 6/7 (85.7%) | 1/1 (100%) |
| MR Locus | 6/7 (85.7%) | 6/793 (0.8%) | 6/6 (100%) | 1/1 (100%) |
| MR-RAPS | 6/7 (85.7%) | 6/793 (0.8%) | 6/6 (100%) | 1/1 (100%) |

